# Seed reserve mobilization and seedling morphology in a bioassay for the detection of genetically modified soybean

**DOI:** 10.1101/2022.07.04.498496

**Authors:** Francisco Cleilson Lopes Costa, Samanda López Peña, Welison Andrade Pereira

**Affiliations:** Universidade Federal de Lavras

**Keywords:** dynamics of seed reserve, *Glycine max*, herbicide

## Abstract

Germination and initial seedling development are physiological processes that determine the establishment of crops. During heterotrophic growth, each seedling develops from the biomass of its seeds. Thus, verifying the potential of genotypes to mobilize reserves under stress has been important. The aim of this study was to investigate how glyphosate affects the mobilization of reserves and seedling morphology. Two tolerant and two herbicide-sensitive cultivars were submitted to germination, seedling length and reserve mobilization tests, including treatments with glyphosate solutions (0, 0.06 and 0.12 %). The hypocotyl / radicle ratio and the efficiency of conversion of reserves to seedlings were also verified. The data were submitted to analysis of variance and test of means. It was observed that the percentage of normal seedlings and length seedlings were affected due to the concentration of the herbicide in the treatments, being the consequences more pronounced for sensitive cultivars; the glyphosate-tolerant genotype and with the best physiological quality mobilized more reserves and was more efficient in converting biomass to seedlings; in morphology, the average length of the seedlings was reduced due to the herbicide, being the roots affected in such a way that they became smaller than the hypocotyls. The herbicide affects the morphology of the seedlings mainly the radicle, and the mobilization of the reserves discriminates the genotypes regarding tolerance to glyphosate.

## Introduction

The germination test (BRASIL, 2009) is one of the first analyses carried out on a seed lot and is performed to assess the germinative potential of the seeds. However, the germination rate observed in the test does not guarantee the establishment of a crop, especially in adverse environmental situations, when the mobilization of seed reserves for seedlings is not satisfactory (OLIVEIRA et al., 2020). In this respect, high vigor seeds are desired by producers because they emerge faster and more uniformly from the soil, accumulate more dry mass in the plants, which are also higher (HENNING et al., 2010).

The initial stages of the seedling development coincide with the period of crop establishment, moment of great apprehension of farmers, aware of the relationship between the formation of the stand and high yield (HENNING et al., 2010). Well-consolidated stand increases soil cover, reduces evaporation of available water, reflecting positively on productivity, improving the competitive power of crops against weeds (DIAS et al., 2011).

According to Mohammadi et al. (2011), the initial weight of the seed, the mobilized fraction of reserves, and the efficiency in converting these reserves to dry matter are relevant components to be evaluated during the heterotrophic growth of the seedlings. During this stage, it is very important to observe how and how much of the seed reserves are mobilized under favorable or unfavorable scenarios for a crop. Studies showed that more vigorous seeds have increased their capacity to mobilize reserves, and consequently, a better initial performance (HENNING et al., 2010; OLIVEIRA et al., 2020).

Crops face adversities from planting to harvest, with germination and seedling emergence being considered the most critical and potentially most sensitive stages to environmental stresses (ALI et al., 2018). Many examples may be presented. In wheat, the seedlings length, the weight of the mobilized reserve, and the fraction of use of this reserve become reduced under water and salt stress (SOLTANI; GHOLIPOOR; ZEINALI, 2006). In soybeans, under water stress, the seedling is impaired, being the development of the hypocotyl is more affected than the radicle (PEREIRA; PEREIRA; DIAS, 2013). The use of desiccants in crops can affect the germination and vigor of harvested soybean seeds, with a negative influence on the mobilization of reserves (DELGADO; COELHO; BUBA, 2015). Copper causes negative effects on cell elongation and cell division in *Vicia sativa* roots and inhibits the mobilization of proteins from cotyledons (MUCCIFORA; BELLANI, 2013). In general, the effects are proportional to the intensity of the stress, which increases the consumption of reserves, without this meaning conversion into dry seedling mass (OLIVEIRA et al., 2020). On the other hand, the pretreatment of soybean seeds with ‘cold plasma’ improved the germination rate, length and dry weight of the seedlings, as well as the mobilization of seed reserves and efficiency in the use of reserves (LING et al., 2014).

Projects on the mobilization of reserves contribute to the understanding of the potential for acclimatization of seedlings (SILVA; AZEVEDO NETO; GHEYI, 2019). Expanding knowledge about the process of mobilizing seed reserves, under the most diverse contexts, can contribute to a better understanding of this important stage. Our research group realized that the availability of genetically modified cultivars for glyphosate tolerance is a good model for studies on the effects of herbicides on the initial development of seedlings. Considered as a model species, our research indeed brings relevant information for soybean and other crops species.

Given the above, the objective of this work was to study the dynamics of soybean seed reserves during the heterotrophic development of the seedlings and their morphology, in an artificial environment simulating the physiological stress that soybean seeds face during germination.

## Materials and Methods

### Water content

To measure the water content (WC) of the studied seed lots, three replicates of 50 seeds (W1) were weighed, taken to the oven at 105 ° C for 24 hours, and weighed again (W2) on a scale with four decimal places. Based on these data, the following formula was applied (SOLTANI; GHOLIPOOR; ZEINALI, 2006).

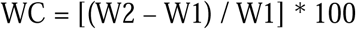

### Germination test

The germination test was carried out according to the recommendations of the Regulation for Seed Analysis (BRASIL, 2009), with adaptations (PEREIRA et al., 2009a), to obtain the first count and the final germination count. Four soybean cultivars were analyzed, 2 sensitive (BRS511, EMGOPA315) and 2 glyphosate-tolerant (TMG1264, P98Y11). The samples were submitted to germination in substrate moistened with water in the control treatment and solutions of the herbicide (0.06 and 0.12%) in the other treatments (PEREIRA et al., 2009a), in the proportion of 2.5 times the weight of dry paper (BRASIL, 2009). A completely randomized design was adopted, in a factorial scheme, composed of 4 genotypes x 3 doses of the herbicide, with 4 replications.

### Seedling length test

The seedling length test was carried out according to (PEREIRA et al., 2009b), with adaptations regarding the moistening of the substrate, with three solutions of the herbicide (0, 0.06 and 0.12% e. a.). A completely randomized design was adopted, in a factorial scheme, 3 cultivars x 3 doses of the herbicide, with 10 repetitions, with each experimental plot consisting of 10 seeds. After the rolls confection, they were packed in plastic bags, with the open end for ventilation, and kept in BOD for 7 days at 25 ° C. At the end of the test, the average lengths of seedling (ALS), hypocotyl (ALH) and radicle (ALR) of each experimental plot were obtained, using a millimeter ruler.

From the average lengths, the proportions of hypocotyl and root of the seedlings in each treatment were known, using the following equations:

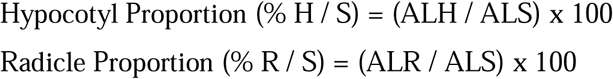

Additionally, the percentages of reduction observed for hypocotyl (RDH) and radicle (RDR) were obtained, due the treatments. For this, the average lengths of the hypocotyl and radicle, under treatment with the herbicide, were compared with those of the experimental control. The following equations were used:

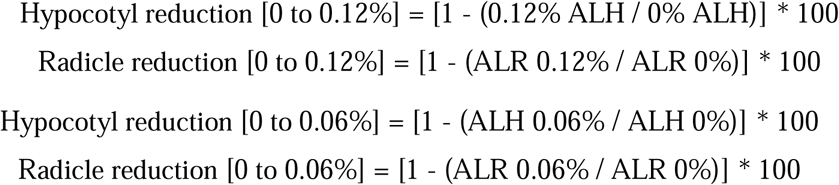

### Reserves Mobilization and Efficiency in Conversion of Reserve to Seedling

Previously, the 120 experimental plots (10 seeds each) separated for the seedling length test were weighed on a precision scale, providing the weight of 10 seeds (W.10) in micrograms. As the water content of each seed lot was previously obtained, it supported the estimation of the dry reserve weight (DW.10) in each experimental plot or in each seed (W.1S) for mobilization from the seeds to seedlings during germination (PEREIRA; PEREIRA; DIAS, 2015).

After measuring the seedlings lengths, the cotyledons, hypocotyls, and radicles of each experimental plot were separated with a stylus and collected in their respective envelopes. Then, these envelopes were kept in an oven at a regulated temperature of 80 °C for 24 hours. From that period, the contents of the envelopes were weighed and the data, in micrograms (μg), used to analyze the mobilization of reserves (PEREIRA; PEREIRA; DIAS, 2015).

Considering the number of seedlings evaluated, the average weight of the dry matter of the hypocotyl (W.1H), radicle (W.1R), seedling (W.1Sd), and cotyledon pair (W.2C) of each experimental plot was obtained. From this information, the first aspect analyzed was the rate of reduction of the seed reserve (% SRR). Given the prior knowledge of the dry weight of the seed (W.1S) and the dry weight of the pair of cotyledons (W.1PC), it was possible to quantify the amount of seed mobilized reserve.

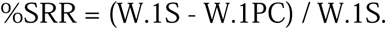

From the knowledge of the dry weight of hypocotyl (W.1H), radicle (W.1R), and seedling (W.1Sd), as well as the reserve that was mobilized from the seeds (SRR = W.1S – W.1PC), it was possible to calculate the proportion of reserve mobilized to the hypocotyl (RMH), radicle (RMR) and seedling (RMS).

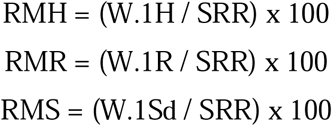

Additionally, we obtained information about the Efficiency of the Conversion of Reserve into Seedling (ERCS). Unlike the variable mobilized reserve for seedling (%SRR), which considers how much of the mobilized reserve has become a seedling, the ECRS variable considers how much of the reserve contained in the seed has become a seedling.

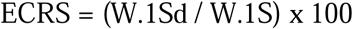

Finally, from the lengths and weights of the seedlings, it was possible to obtain the linear weights of hypocotyl (LWH), radicle (LWR) and seedling (LWS) in μg / cm.

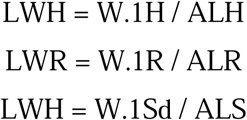

### Experimental design and data analysis

The tests were performed in a completely randomized design, in a factorial scheme. The number of repetitions and the size of the experimental plots varied between experiments. The data were subjected to analysis of variance and means compared by the Scott-Knott test, at the level of 5% probability.

## Results and Discussion

### Germination test

Analyzing the first and the final counts of germination data (Table 1), it was notable that both herbicide solutions (0.06, and 0.12 %) inhibited the development of normal seedlings by the genotypes, like observed also by Bervald et al. (2010) and Pereira et al., (2018). In the case of glyphosate-tolerant cultivars, the highest concentration of the herbicide (0.12 %) affected the first and the final counts of germination, as observed by Pereira et al., (2018), however, the glyphosate-tolerant seedlings developed secondary roots, while the sensitive ones did not (Figure 1). The absence of secondary roots in glyphosate-sensitive seedlings is a striking feature of this bioassay (BERVALD et al., 2010; MELO; FAGIOLI; SÁ, 2013; PÁDUA et al., 2012). According to Albrecht et al. (2014), the physiological quality and the vigor of the lots fall due to the concentration of the herbicide.

**Table 1.** First count and final count of the germination of four soybean cultivars submitted to germination test on moistened substrate with glyphosate solutions.

| Dosage | First Count of Germination (%) |  |  |  | Final Count of Germination (%) |  |  |  |
| --- | --- | --- | --- | --- | --- | --- | --- | --- |
|  | Emgopa<br>315 | UFUS<br>7415 | P98Y11 | TMG<br>1264 | Emgopa<br>315 | UFUS<br>7415 | P98Y11 | TMG<br>1264 |
| 0.00 % | 76.5 Ba | 52.0 Ca | 54.0 Ca | 91.5 Aa | 78.0 Ba | 56.5 Ca | 59.5 Ca | 94.0 Aa |
| 0.06 % | 0.5 Cb | 0.0 Cb | 58.0 Ba | 80.5 Ab | 1.5 Cb | 0.1 Cb | 60.5 Ba | 94.0 Aa |
| 0.12 % | 0.0 Bb | 0.0 Bb | 5.5 Bb | 62.0 Ac | 0.5 Cb | 0.1 Cb | 27.5 Bb | 78.0 Ab |
| Mean |  | 37.4 |  |  |  | 45.4 |  |  |
| CV(%) |  | 26.7 |  |  |  | 20.4 |  |  |

\* Averages followed by the same letter, uppercase in the horizontal and lowercase in the vertical, do not differ each other, at the level of 5 % probability.

**Figure 1.**
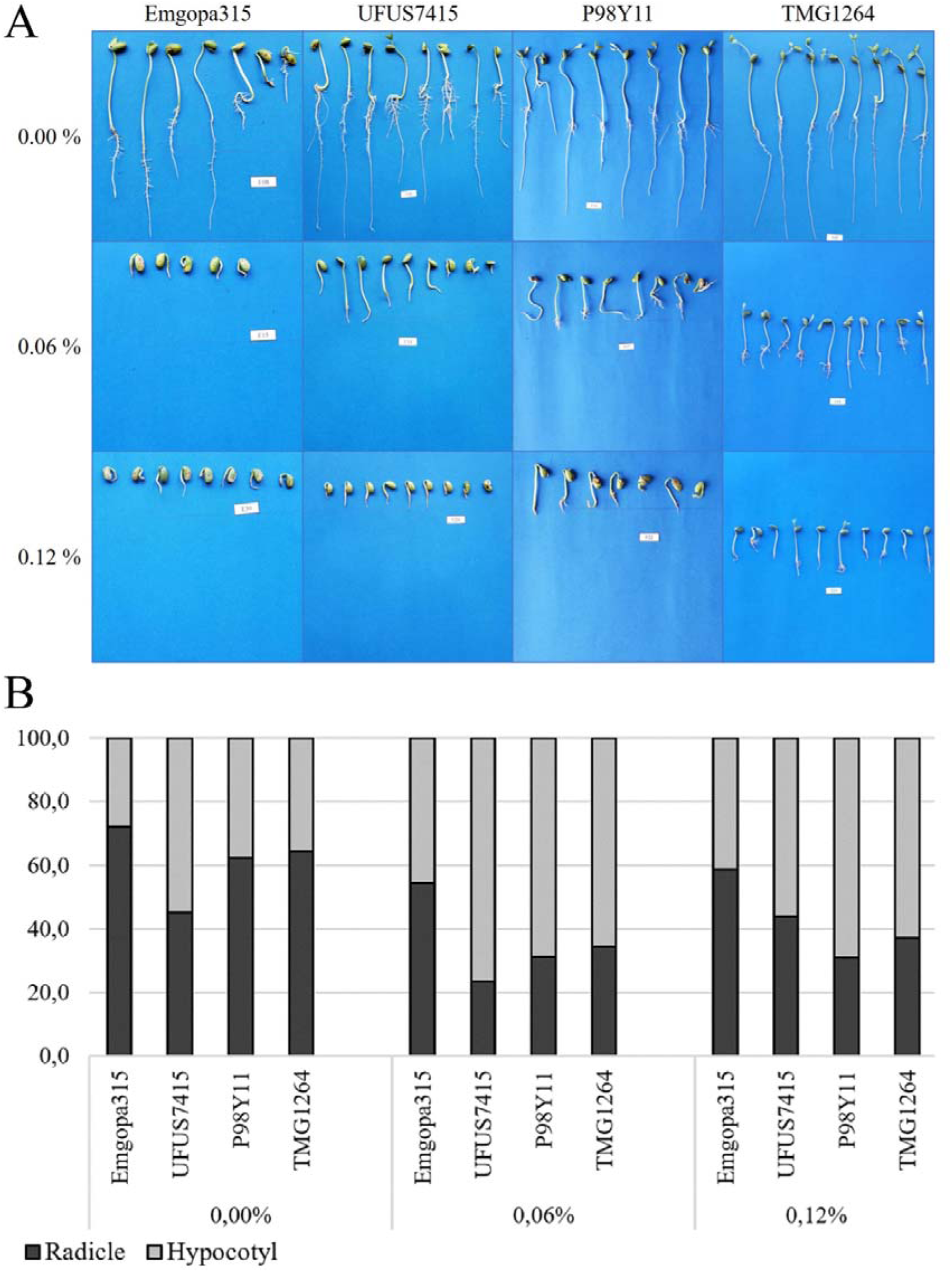
Seedling morphology (A), hypocotyl and radicle proportions (B) of Emgopa315, UFUS7415, P98Y11, and TMG1264 cultivars, submitted to germination in substrate moistened with different glyphosate solutions (0, 0.06 and 0.12 %).

In a work assessing the efficiency of toothpeeks mostened with glyphosate to diferentiate tolerant and non-tolerant soybean cultivars, Silva et al. (2015) stated that the injured plants of transgenic cultivars keeped growing the hypocotyl (main stem) and developed new lateral branches instead of new lateral radicles like in our research. This differenc may be explained because their role plants were not exposed to glyphosate at the initial growing stages, then the glyphosate followed the xylem flux to the aerial parts of the seedlings. Differently, our seedlings were completely exposed to the glyphosate solution treatments, then we can affirm the radicule tissues are more sensitive to glyphosate, since no secondary branches were observed in our results.

### Seedling length test

The herbicide treatments reduced the lengths of the hypocotyl (ALH), radicle (ALR), and seedling (ALS) (Table 2). In the herbicide treatments, the secondary roots are absent or incipient in sensitive seedlings, with an aspect of blocked development. Cultivars that carry the CP4 EPSPS gene are glyphosate-tolerant (NAKATANI et al., 2014). When these cultivars are submitted to germination in substrate moistened with glyphosate solutions, secondary roots are present, but in length, number, and volume smaller than in the control treatment, as observed by Pereira et al., (2018); in fact, the herbicide affects the development of sensitive and tolerant seedlings at different levels of intensity.

**Table 2.** Average length of hypocotyl, radicle and seedling of four soybean cultivars treated with three (0, 0.06, and 0.12 %) doses of glyphosate.

| Average length of the hypocotyl - ALH (cm) |  |  |  |  |
| --- | --- | --- | --- | --- |
|  | Emgopa 315 | UFUS 7415 | P98Y11 | TMG 1264 |
| 0.00 % | 5.12 Ca | 7.89 Aa | 6.79 Ba | 7.54 Aa |
| 0.06 % | 1.66 Cb | 3.65 Bb | 5.09 Ab | 5.67 Ab |
| 0.12 % | 1.22 Bb | 1.42 Bc | 3.83 Ac | 3.87 Ac |
| Mean | 4,5 |  |  |  |
| CV(%) | 18,8 |  |  |  |
| Average length of the radicle - ALR (cm) |  |  |  |  |
|  | Emgopa 315 | UFUS 7415 | P98Y11 | TMG 1264 |
| 0.00 % | 13.2 Aa | 6.48 Ca | 11.3 Ba | 13.5 Aa |
| 0.06 % | 1.91 Bb | 1.15 Bb | 2.28 Ab | 3.03 Ab |
| 0.12 % | 1.74 Ab | 1.08 Ab | 1.68 Ab | 2.33 Ab |
| Mean | 4,9 |  |  |  |
| CV (%) | 31,3 |  |  |  |
| Average length of the seedling - ALS (cm) |  |  |  |  |
|  | Emgopa 315 | UFUS 7415 | P98Y11 | TMG 1264 |
| 0.00 % | 18.3 Ba | 14.4 Ca | 18.1 Ba | 21.0 Aa |
| 0.06 % | 3.56 Bb | 4.80 Bb | 7.38 Ab | 8.71 Ab |
| 0.12 % | 2.96 Bb | 2.50 Bc | 5.51 Ac | 6.19 Ac |
| Mean | 9,4 |  |  |  |
| CV (%) | 17,4 |  |  |  |

\* Averages followed by the same letter, uppercase in the horizontal and lowercase in the vertical, do not differ, at the level of 5 % probability.

According to Funguetto et al. (2004); Heinz; Viegas Neto and Valente (2011); Pereira et al. 2018; and Tillmann and West (2004) the length reductions are more striking for glyphosate-sensitive genotypes. In our study, this was a notable occurrence.

Once it was verified the average lengths (ALH, ALR, and ALP) are reduced, it was questioned whether treatments with the herbicide would proportionally affect hypocotyls and radicles (Figure 1). It was observed that at a dose of 0 %, the radicle length was about 2/3 of the seedling length. But, in the doses of 0.06 and 0.12 %, the seedlings presented hypocotyls in greater proportion. There was an inversion in the seedling architecture after the treatment with glyphosate. Heinz; Viegas Neto and Valente (2011) also noticed a reduction in the radicle length. In our study, the hypocotyl becomes larger than the primary root when the seedlings were treated with the herbicide. In the Chinese cabbage seed, tetracycline, whose mechanism of action occurs at the ribosomes level, blocked the mobilization of protein bodies in cotyledons during radicle elongation (LUO et al., 2020). We believe that the explanation is plausible, since glyphosate affects protein synthesis by blocking the EPSPS enzyme (NAKATANI et al., 2014).

For these results, it is possible to assume that the development of the aerial part would become a priority during the development with glyphosate. This result is interesting, considering that was observed in other studies that the seed lots with contrasting physiological quality would have no difference for dry matter weight in the hypocotyl, but for radicle (OLIVEIRA et al., 2020). Thus, other metabolic pathways can be activated in these conditions.

To better investigate this observation, the average length of the hypocotyl (ALH) and radicle (ALR) in herbicide treatments (0.06 and 0.12 %) were divided by the respective values found in the control treatment (0 %), and the results obtained in percentages.

About the hypocotyl, the rates of reductions in the genotypes Emgopa315, UFUS7415, P98Y11, and TMG1264 were 68, 53, 25, and 25%, respectively, in the 0.06% treatment. In the 0.12% treatment, these rates increase to 76, 82, 44, and 49%, respectively. We observe that the reductions were more intense for glyphosate-sensitive cultivars. The herbicide has an inhibitory effect on the organogenesis of sensitive seedlings (BERVALD et al., 2010).

In the case of radicles, unlike the hypocotyl, all genotypes were similarly affected, both in the treatment of 0.06% and 0.12%, reinforcing the hypothesis that glyphosate affects radicle development more than the hypocotyl, also seen in soybean by Tillmann and West (2004) and Heinz et al. (2011), and in pea by Mondal et al. (2017).

These results show that the radicles are harmed by the herbicide in a drastic way, while the hypocotyl becomes a priority in seedling development. However, even though the hypocotyl has priority, sensitive cultivars do not develop it well, because of EPSPS inhibition.

### Reserve mobilization

The weights of the hypocotyl (W.1H), radicle (W.1R), and seedling (W.1Sd) of the genotypes were reduced due of the herbicide treatments (Table 3). The glyphosate-tolerant genotypes showed more dry matter than the glyphosate-sensitive genotypes. This is an interesting result because glyphosate-sensitive and glyphosate-tolerant cultivars could not be differentiated by ALR, but by W.1R; this was possible because the roots of tolerant seedlings are heavier (TORRES et al., 2003). In fact, the average root length does not include secondary roots, developed only by herbicide-tolerant cultivars, and absent or incipient in sensitive genotypes (FUNGUETTO et al., 2004; MELO; FAGIOLI; SÁ, 2013; PÁDUA et al., 2012).

**Table 3.** Dry matter weigh of hypocotyl, radicle and seedling of four soybean cultivars treated with three (0, 0.06, and 0.12 %) doses of glyphosate.

| Dry matter weigh of hypocotyl – P.1H (µg) |  |  |  |  |
| --- | --- | --- | --- | --- |
| Dosage | Emgopa 315 | UFUS 7415 | P98Y11 | TMG 1264 |
| 0.00 % | 14.40 Ba | 25.30 Aa | 26.7 Aa | 27.9 Aa |
| 0.06 % | 7.23 Cb | 15.40 Bb | 21.5 Ab | 23.1 Ab |
| 0.12 % | 5.14 Cc | 8.85 Bc | 18.8 Ac | 17.8 Ac |
| Mean | 17.7 |  |  |  |
| CV (%) | 12.4 |  |  |  |
| Dry matter weigh of radicle – P.1R (µg) |  |  |  |  |
| Dosage | Emgopa 315 | UFUS 7415 | P98Y11 | TMG 1264 |
| 0.00 % | 4.70 Ca | 10.80 Aa | 9.56 Ba | 10.70 Aa |
| 0.06 % | 1.95 Cb | 4.04 Bb | 5.33 Ab | 6.16 Ab |
| 0.12 % | 1.68 Db | 3.47 Cb | 4.82 Bb | 5.91 Ab |
| Mean | 5.8 |  |  |  |
| CV (%) | 20.3 |  |  |  |
| Dry matter weigh of seedling – P.2C (µg) |  |  |  |  |
| Dosage | Emgopa 315 | UFUS 7415 | P98Y11 | TMG 1264 |
| 0.00 % | 19.10 Ba | 36.1 Aa | 36.2 Aa | 38.6 Aa |
| 0.06 % | 9.18 Cb | 19.4 Bb | 26.8 Ab | 29.2 Ab |
| 0.12 % | 6.82 Cb | 12.3 Bc | 23.6 Ac | 23.7 Ac |
| Mean | 23.4 |  |  |  |
| CV (%) | 11.8 |  |  |  |

\* Averages followed by the same letter, uppercase in the horizontal and lowercase in the vertical, do not differ, at the level of 5 % probability.

Regarding the contrast between the dry matter contained in the seeds and the dry matter of the seedlings, it was possible to estimate the percentage of dry matter mobilized for the hypocotyls, roots, and seedlings (Table 4).

**Table 4.**
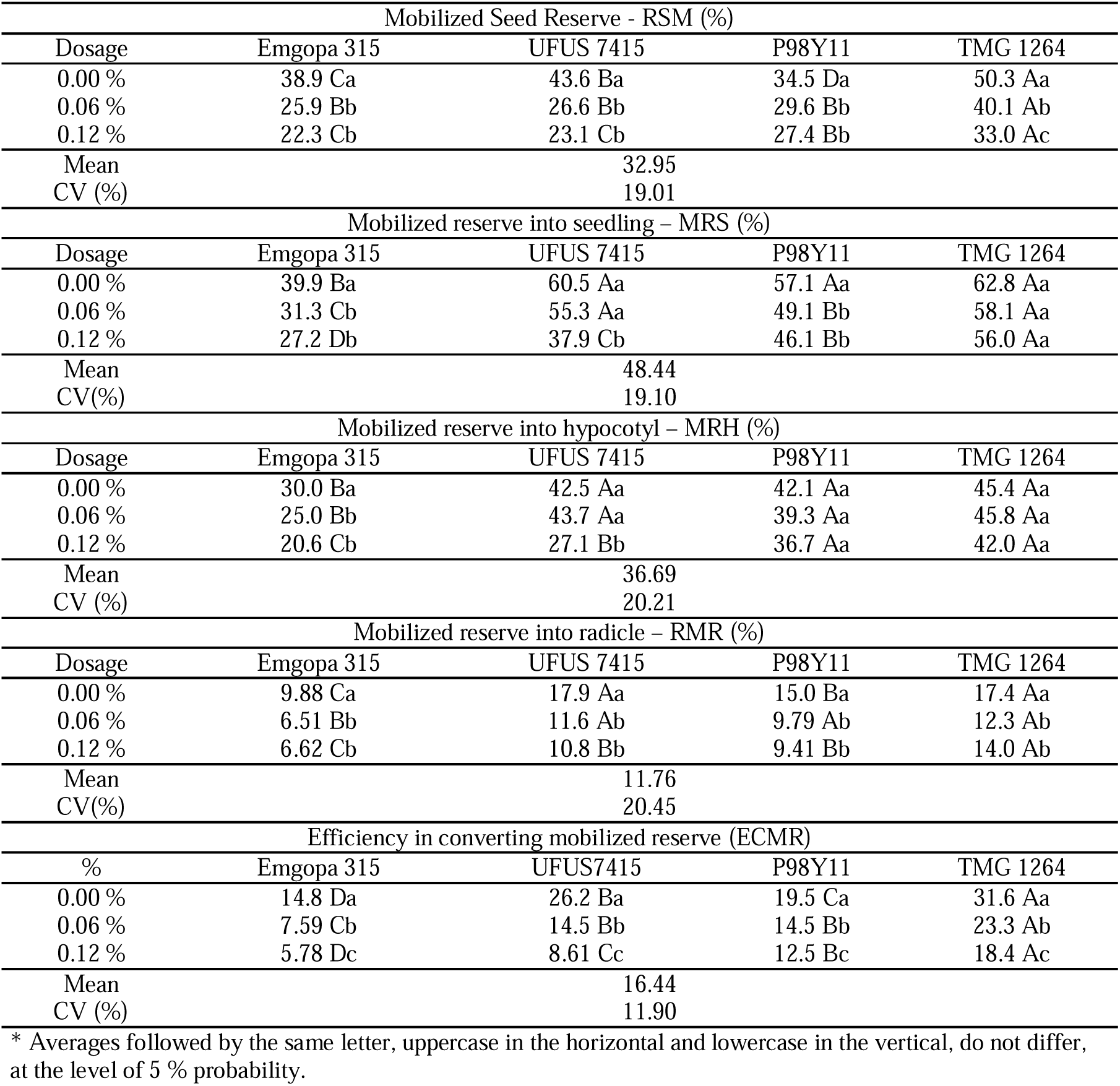
Mobilization of reserves for hypocotyl, radicle and seedling, and Efficiency in converting mobilized reserves (ECMR) of soybean cultivars treated with three (0, 0.06 and 0.12 %) doses of glyphosate.

| Mobilized Seed Reserve - RSM (%) |  |  |  |  |
| --- | --- | --- | --- | --- |
| Dosage | Emgopa 315 | UFUS 7415 | P98Y11 | TMG 1264 |
| 0.00 % | 38.9 Ca | 43.6 Ba | 34.5 Da | 50.3 Aa |
| 0.06 % | 25.9 Bb | 26.6 Bb | 29.6 Bb | 40.1 Ab |
| 0.12 % | 22.3 Cb | 23.1 Cb | 27.4 Bb | 33.0 Ac |
| Mean |  | 32.95 |  |  |
| CV (%) |  | 19.01 |  |  |
| Mobilized reserve into seedling – MRS (%) |  |  |  |  |
| Dosage | Emgopa 315 | UFUS 7415 | P98Y11 | TMG 1264 |
| 0.00 % | 39.9 Ba | 60.5 Aa | 57.1 Aa | 62.8 Aa |
| 0.06 % | 31.3 Cb | 55.3 Aa | 49.1 Bb | 58.1 Aa |
| 0.12 % | 27.2 Db | 37.9 Cb | 46.1 Bb | 56.0 Aa |
| Mean |  | 48.44 |  |  |
| CV(%) |  | 19.10 |  |  |
| Mobilized reserve into hypocotyl – MRH (%) |  |  |  |  |
| Dosage | Emgopa 315 | UFUS 7415 | P98Y11 | TMG 1264 |
| 0.00 % | 30.0 Ba | 42.5 Aa | 42.1 Aa | 45.4 Aa |
| 0.06 % | 25.0 Bb | 43.7 Aa | 39.3 Aa | 45.8 Aa |
| 0.12 % | 20.6 Cb | 27.1 Bb | 36.7 Aa | 42.0 Aa |
| Mean |  | 36.69 |  |  |
| CV (%) |  | 20.21 |  |  |
| Mobilized reserve into radicle – RMR (%) |  |  |  |  |
| Dosage | Emgopa 315 | UFUS 7415 | P98Y11 | TMG 1264 |
| 0.00 % | 9.88 Ca | 17.9 Aa | 15.0 Ba | 17.4 Aa |
| 0.06 % | 6.51 Bb | 11.6 Ab | 9.79 Ab | 12.3 Ab |
| 0.12 % | 6.62 Cb | 10.8 Bb | 9.41 Bb | 14.0 Ab |
| Mean |  | 11.76 |  |  |
| CV(%) |  | 20.45 |  |  |
| Efficiency in converting mobilized reserve (ECMR) |  |  |  |  |
| % | Emgopa 315 | UFUS7415 | P98Y11 | TMG 1264 |
| 0.00 % | 14.8 Da | 26.2 Ba | 19.5 Ca | 31.6 Aa |
| 0.06 % | 7.59 Cb | 14.5 Bb | 14.5 Bb | 23.3 Ab |
| 0.12 % | 5.78 Dc | 8.61 Cc | 12.5 Bc | 18.4 Ac |
| Mean |  | 16.44 |  |  |
| CV (%) |  | 11.90 |  |  |

\* Averages followed by the same letter, uppercase in the horizontal and lowercase in the vertical, do not differ, at the level of 5 % probability.

First, about the hypocotyl, it was noted that treatments with the herbicide reduced the reserve rate mobilized in cultivars herbicide-sensitive, Emgopa315 (treatments 0.06 and 0.12%) and UFUS7415 (treatment 0.12%); however, for the tolerant genotypes (P98Y11 and TMG1264), there was no reduction in the mobilized reserve due to the herbicide. Given these results, the development of the hypocotyl, in fact, is different for cultivars sensitive and tolerant to glyphosate (HEINZ; VIEGAS NETO; VALENTE, 2011; TILLMANN; WEST, 2004).

For the radicle, regardless of the genotype and the concentration of the herbicide, the mobilization of reserves was reduced, which corroborates the observation that the radicles of the four genotypes are affected equally. Therefore, in the case of the radicle, which is more affected than the hypocotyl in treatment with the herbicide (TILLMANN; WEST, 2004), neither the length nor the weight or the reserve mobilization efficiently differentiated the cultivars in terms of tolerance or sensitivity to the herbicide. As presented by Torres et al. (2003), we also consider root development as a marker feature to differentiate genotypes in relation to glyphosate tolerance, especially with regard to the development of secondary roots.

In the case of seedlings, the effects varied. There were differences between classes of genotypes, herbicide-tolerant and herbicide-sensitive, as well as between sensitive genotypes (Emgopa315 and UFUS7415) and between tolerant genotypes (P98Y11 and TMG1264). Except for TMG1264, the treatments with the herbicide (0.06 and/or 0.12 %) reduced the reserve mobilized into the seedlings. Possibly, it is a result associated with the better quality of the seed lot (ANDRADE; COELHO; PADILHA, 2019). Anyway, the accumulation of dry matter in seedlings from stressed seeds is less than that of non-stressed seeds (SILVA; AZEVEDO NETO; GHEYI, 2019).

### Efficiency of Conversion of Seed Reserve to Seedling

Regarding the percentage of mobilized reserves of the cotyledons, the herbicide affected all genotypes at different levels (Table 4). Notably, the glyphosate-tolerant genotype and with better physiological quality, TMG1264, was the one that most mobilized reserves of its cotyledons and presented the greatest efficiency in converting seed reserves into seedlings. Further studies should assess whether this is a particularity of the genotype or an issue of initial physiological quality, but there are literatures reporting the better quality lots result in seedlings with more dry matter (ANDRADE; COELHO; PADILHA, 2019; OLIVEIRA et al., 2020). Anyway, a focused investigation on the subject is necessary, because if there is an important genetic factor controlling this trait, genetic improvement can be applied.

Notable characteristics of the bioassay in this study were the reduction in the length and thickening of the hypocotyl-radicle axis of the seedlings. To verify this, the linear weights (ug / cm) of the hypocotyl (LWH), root (LWR) and seedling (LWS) were estimated (Table 5). We observed that the genotypes increased their linear weights due to treatments with herbicides. Under these conditions, the seedlings concentrate more reserves, thickening the hypocotyl-radicle axis, as also seen by Funguetto et al. (2004). This result corroborates with the herbicide effect in reducing the seedlings length. Considering that both the seedlings length and the dry matter weight of the seedlings were affected by treatments with the herbicide, the linear weight showed that the length was more affected than the weight since the linear weight increased due to the herbicide concentration in all treatments.

**Table 5.**
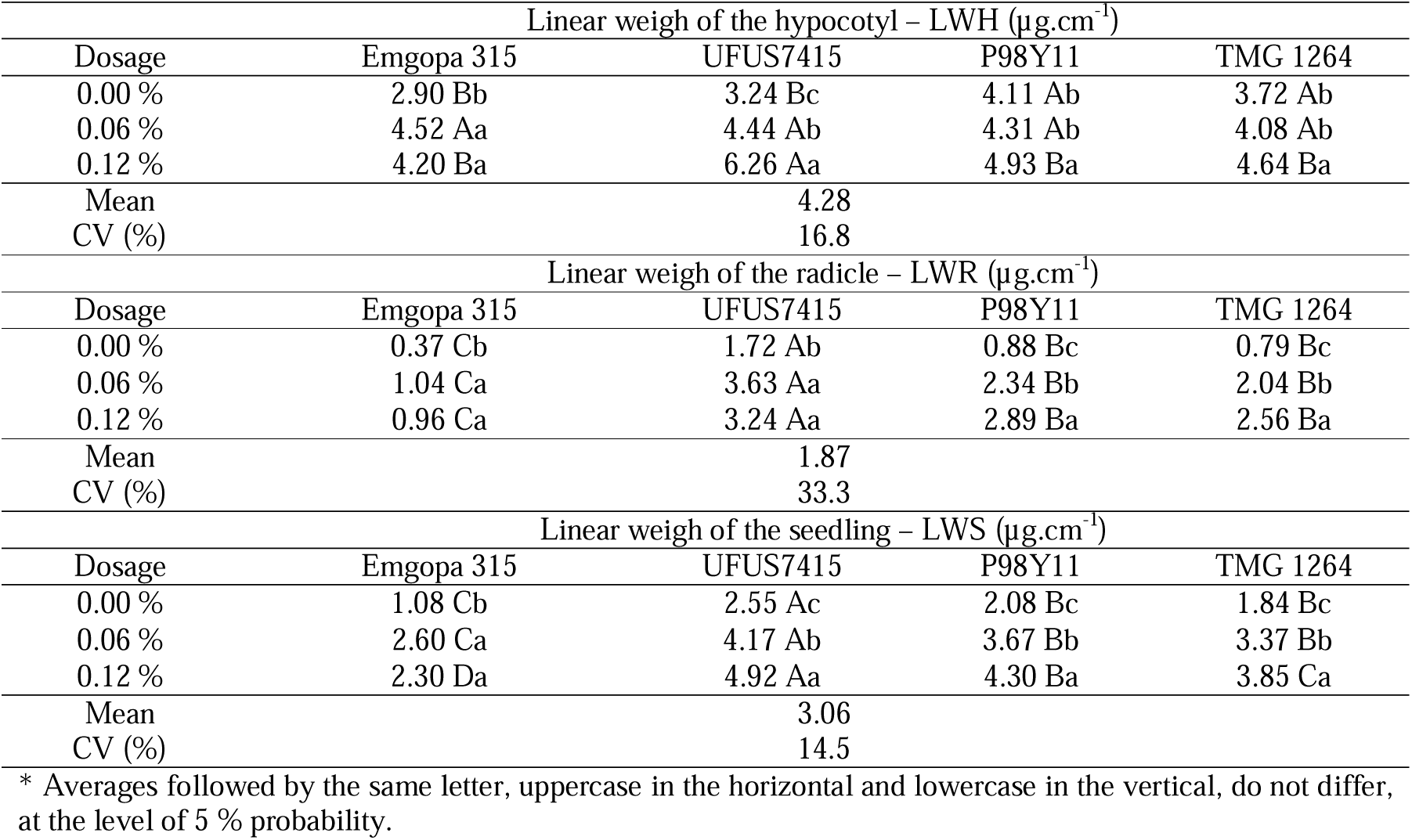
Hypocotyl, radicle and seedling linear weigh of four soybean cultivars treated with three (0, 0.06 and 0.12 %) doses of glyphosate.

It was possible to observe that some responses of the genotypes vary in relation to the herbicide, while others are common to all. In fact, herbicide-tolerant cultivars form normal seedlings under treatment with glyphosate but show sensitivity at certain concentrations. Length, weight and mobilization of reserves are fundamental characteristics of the initial phase of seedling development and can be considered in breeding programs. The identification of QTLs associated with the use of seed reserves in selected RILs (CHENG et al., 2013), for example, provided a glimpse into the use of marker-assisted selection to improve the mobilization of rice reserves.

## Conclusions

1. The herbicide affects the length, weight, and reserves mobilization of the evaluated genotypes, but in a more striking way those that are glyphosate-sensitive.
2. During germination, the herbicide affects the radicle more than the seedling hypocotyl, making it proportionally larger than the radicle, changing the seedling morphology.
3. The mobilization of reserves discriminates genotypes regarding tolerance to the herbicide, reinforcing and complementing the seedlings length analyzes.

## Acknowledgements

The Coordenação de Aperfeiçoamento de Pessoal de Nível Superior (CAPES), for providing the scholarships to the authors FCLC and SLP.

## References

Albrecht, A. J. P. et al. Behavior of RR soybeans subjected to different formulations and rates of glyphosate in the reproductive period. Planta Daninha, v. 32, n. 4, p. 851–859, 2014. DOI: 10.1590/S0100-83582014000400020

Ali, Q. et al. Assessment of drought tolerance in mung bean cultivars/lines as depicted by the activities of germination enzymes, seedling’s antioxidative potential and nutrient acquisition. Archives of Agronomy and Soil Science, v. 64, n. 1, p. 84–102, 2 jan. 2018. DOI: 10.1080/03650340.2017.1335393

Andrade, G. C. DE; Coelho, C. M. M.; Padilha, M. S. Seed reserves reduction rate and reserves mobilization to the seedling explain the vigour of maize seeds. Journal of Seed Science, v. 41, n. 4, p. 488–497, 5 dez. 2019. DOI: 10.1590/2317-1545v41n4227354

Bervald, C. M. P. et al. Desempenho fisiológico de sementes de soja de cultivares convencional e transgênica submetidas ao glifosato. Revista Brasileira de Sementes, v. 32, n. 2, p. 09–18, jun. 2010. DOI: 10.1590/S0101-31222010000200001

BRASIL. Regras para análise de sementes. Brasília: Ministério da Agricultura, Pecuária e Abastecimento, 2009.

Cheng, X. et al. Dynamic Quantitative Trait Loci Analysis of Seed Reserve Utilization during Three Germination Stages in Rice. PLOS ONE, v. 8, n. 11, p. e80002, 11 nov. 2013. DOI: 10.1371/journal.pone.0080002

Delgado, C. M. L.; Coelho, C. M. M. DE; Buba, G. P. Mobilization of reserves and vigor of soybean seeds under desiccation with glufosinate ammonium. Journal of Seed Science, v. 37, n. 2, p. 154–161, 14 jul. 2015. DOI: 10.1590/2317-1545v37n2148445

Dias, M. A. N. et al. Direct effects of soybean seed vigor on weed competition. Revista Brasileira de Sementes, v. 33, n. 2, p. 346–351, 2011. DOI: 10.1590/S0101-31222011000200017

Funguetto, C. I. et al. Detecção de sementes de soja geneticamente modificada tolerante ao herbicida glifosato. Revista Brasileira de Sementes, v. 26, n. 1, p. 130–138, 2004. DOI: 10.1590/S0101-31222004000100020

Heinz, R.; Viegas Neto, A. L.; Valente, T. DE O. Detecção de sementes de soja geneticamente modificada por meio de teste de germinação. Agrarian, v. 4, n. 11, p. 20–26, 2011. DOI: 10.1590/S0006-87052010000300026

Henning, F. A. et al. Composição química e mobilização de reservas em sementes de soja de alto e baixo vigor. Bragantia, v. 69, n. 3, p. 727–734, 2010. DOI: 10.1590/S0006-87052010000300026

Ling, L. et al. Effects of cold plasma treatment on seed germination and seedling growth of soybean. Scientific Reports 2014 4:1, v. 4, n. 1, p. 1–7, 31 jul. 2014. DOI: 10.1038/srep05859

Luo, Y. et al. Tetracycline stress disturbs the mobilization of protein bodies in seed storage reserves during radicle elongation after seed germination. Environmental Science and Pollution Research 2020 27:33, v. 27, n. 33, p. 42150–42157, 26 ago. 2020. DOI: 10.1007/s11356-020-10569-7

Melo, L. F.; Fagioli, M.; Sá, M. E. Alternative methods for detecting soybean seeds genetically modified for resistance to herbicide glyphosate. Journal of Seed Science, v. 35, n. 3, p. 381–386, 2013. DOI: 10.1590/S2317-15372013000300016

Mohammadi, H. et al. Effects of seed aging on subsequent seed reserve utilization and seedling growth in soybean. International Journal of Plant Production, v. 5, n. 1, p. 65–70, 2011.

Mondal, S. et al. Phytotoxicity of glyphosate in the germination of Pisum sativum and its effect on germinated seedlings. Environmental Health and Toxicology, v. 32, p. e2017011, 16 ago. 2017. DOI: 10.5620/eht.e2017011

Muccifora, S.; Bellani, L. M. Effects of copper on germination and reserve mobilization in Vicia sativa L. seeds. Environmental Pollution, v. 179, p. 68–74, 1 ago. 2013. DOI: 10.1016/j.envpol.2013.03.061

Nakatani, A. S. et al. Effects of the glyphosate-resistance gene and of herbicides applied to the soybean crop on soil microbial biomass and enzymes. Field Crops Research, v. 162, p. 20–29, 1 jun. 2014. DOI: 10.1016/j.fcr.2014.03.010

Oliveira, T. F. et al. Reserve mobilization in soybean seeds under water restriction after storage. Journal of Seed Science, v. 42, p. 1–8, 14 ago. 2020. DOI: 10.1590/2317-1545v42231384

Pádua, G. P. DE et al. Detection of adventitious presence of genetically modified seeds in lots of non transgenic soybean. Revista Brasileira de Sementes, v. 34, n. 4, p. 573–579, 2012. DOI: 10.1590/S0101-31222012000400007

Pereira, W. A. et al. Ajuste de metodologias para a identificação de cultivares de soja quanto à tolerância ao glifosato. Revista Brasileira de Sementes, v. 31, n. 4, p. 133–144, 2009. DOI: 10.1590/S0101-31222009000400016

Pereira, W. A. et al. Influência da disposição, número e tamanho das sementes no teste de comprimento de plântulas de soja. Revista Brasileira de Sementes, v. 31, n. 1, p. 113–121, 2009b. DOI: 10.1590/S0101-31222009000100013

Pereira, W. A. et al. Performance of transgenic and conventional soybean plants subjected to bioassay for detection of glyphosate tolerant seeds. Crop Breeding and Applied Biotechnology, v. 18, n. 1, p. 39–46, 2018. DOI: 10.1590/1984-70332018v18n1a6

Pereira, W. A.; Pereira, S. M. A.; Dias, D. C. F. Dos S. Influence of seed size and water restriction on germination of soybean seeds and on early development of seedlings. Journal of Seed Science, v. 35, n. 3, p. 316–322, 2013.

Pereira, W. A.; Pereira, S. M. A.; Dias, D. C. F. Dos S. Dynamics of reserves of soybean seeds during the development of seedlings of different commercial cultivars. Journal of Seed Science, v. 37, n. 1, p. 63–69, 2015.

Silva, F. M. DA et al. Toothpick test: a methodology for the detection of RR soybean plants. Revista Ciência Agronômica, v. 46, n. 2, 2015. DOI: 10.5935/1806-6690.20150024

Silva, P. C. C.; Azevedo Neto, A. D. DE; Gheyi, H. R. Mobilization of seed reserves pretreated with H2O2 during germination and establishment of sunflower seedlings under salinity. Journal of Plant Nutrition, v. 42, n. 18, p. 2388–2394, 2019. DOI: 10.1080/01904167.2019.1659349

Soltani, A.; Gholipoor, M.; Zeinali, E. Seed reserve utilization and seedling growth of wheat as affected by drought and salinity. Environmental and Experimental Botany, v. 55, n. 1, p. 195–200, 2006. DOI: 10.1016/j.envexpbot.2004.10.012

Tillmann, M. Â. A.; West, S. Identification of geneticaly modified soybean seeds resistant to glyphosate. Scientia Agricola, v. 61, n. 3, p. 336–341, 2004. DOI: 10.1590/S0103-90162004000300017

Torres, A. C. et al. Bioassay for detection of transgenic soybean seeds tolerant to glyphosate. Pesquisa Agropecuária Brasileira, v. 38, n. 9, set. 2003. DOI: 10.1590/S0100-204X2003000900005

